# Evolution of multi-partner symbiotic systems in the tribe Cerataphidini: genome reduction of Buchnera and frequent turnover of companion symbionts

**DOI:** 10.1101/2025.07.09.664008

**Authors:** Shunta Yorimoto, Mitsuru Hattori, Tomonari Nozaki, Shuji Shigenobu

## Abstract

How multi-partner symbiotic systems originate and diversify within a lineage remains poorly understood. To address this, we investigated the evolutionary dynamics of bacterial symbionts across eight species of the aphid subfamily Hormaphidinae, with detailed molecular, genomic, and microscopic analyses focused on two Cerataphidini species, *Ceratovacuna nekoashi* and *Pseudoregma panicola*. Our 16S ribosomal RNA gene amplicon sequencing analysis revealed that Cerataphidini species consistently harbor companion symbionts alongside *Buchnera*, whereas the examined Hormaphidini and Nipponaphidini species harbor only *Buchnera*. Notably, *Arsenophonus* in *C. nekoashi* and *C. japonica* have distinct phylogenetic origins, and *P. panicola* has acquired *Symbiopectobacterium* instead of *Arsenophonus*. Microscopic analyses demonstrated that these companion symbionts are maternally transmitted and occupy distinct cell types from those harboring *Buchnera*. Genome sequencing revealed extreme reduction in *Buchnera* genomes to ∼0.4 Mbp in both *C. nekoashi* and *P. panicola*, comparable to *C. japonica* but significantly smaller than *Buchnera* genomes of ∼0.6 Mbp in mono-symbioses. The companion symbiont genomes retain complete riboflavin biosynthesis pathways absent in *Buchnera*. Our findings suggest that *Buchnera* genome reduction and companion symbiont acquisition occurred in the common ancestor of Cerataphidini, followed by multiple companion symbiont replacements, demonstrating that ancient obligate symbionts can remain evolutionarily stable while companion symbionts undergo frequent turnover.

**Importance:** Many insects depend on multiple bacterial symbionts to survive on nutritionally limited diets, yet how such multi-partner symbiotic systems originate and change over evolutionary time is poorly understood. By examining the aphid subfamily Hormaphidinae, particularly *Ceratovacuna nekoashi* and *Pseudoregma panicola*, we show that a dramatic reduction in the genome of the ancient symbiont *Buchnera* and the acquisition of a companion symbiont occurred together in the common ancestor of the tribe Cerataphidini. Strikingly, while *Buchnera* has been stably maintained across all examined species, the companion symbiont has been repeatedly replaced by phylogenetically diverse bacteria, even among closely related species. These findings reveal an evolutionary asymmetry between ancient and recently acquired symbionts and suggest that companion symbiont turnover is a common consequence in insects harboring genomically degraded obligate symbionts.

## Introduction

Symbiotic relationships between animals and microorganisms are ubiquitous in nature, with many insect species relying on inherited endosymbiotic bacteria for essential nutrients (1, 2). These associations range from single-partner to complex multi-partner symbiotic systems. Insects have evolved specialized host cells known as bacteriocytes to harbor bacterial symbionts, and these symbionts are vertically transmitted to progeny (3). However, this intracellular lifestyle leads to symbiont genome degradation and accumulation of deleterious mutations in the host-restricted symbionts (4–6). The host insects often acquire additional symbionts, forming co-obligate partnerships where the companion symbionts complement the functions lost in the primary symbionts (7, 8). These multi-partner symbiotic systems have evolved multiple times in hemipteran insects (8, 9). Within this order, aphids (Hemiptera: Aphididae) exhibit remarkable diversity in symbiotic associations, ranging from ancient mono-symbiotic to recently evolved co-obligate symbiotic systems (10). This variation across aphid lineages provides opportunities to investigate how multi-partner symbioses arise and diversify.

Almost all aphid species harbor *Buchnera aphidicola*, an obligate mutualistic symbiont that provides essential nutrients lacking in their sole diet of plant phloem sap (11, 12). The genomes of *Buchnera* have undergone drastic reduction to approximately 0.6 Mbp with around 600 protein-coding genes, while retaining genes crucial for synthesizing the essential nutrients (13–15). *Buchnera* resides within bacteriocytes and is vertically transmitted to progeny (16, 17). In addition to *Buchnera*, many aphid lineages have acquired secondary symbionts, which may be either facultative or co-obligate. Facultative symbionts can enhance host fitness under specific environmental conditions (18), while co-obligate symbionts have become essential by complementing degraded functions in the ancient obligate symbiont. In these co-obligate symbioses, such as those within the subfamilies Lachninae and Chaitophorinae, the genomes of *Buchnera* have undergone even more extensive reduction, shrinking to approximately 0.4 Mbp with around 400 protein-coding genes (19–22). These reduced genomes lack genes for synthesizing some essential amino acids and B vitamins, notably riboflavin. The metabolic deficiencies in *Buchnera* are complemented by additional co-obligate symbiont, *Serratia symbiotica*, which resides in bacteriocytes distinct from those harboring *Buchnera* (20–23).

Recent research has revealed a novel co-obligate endosymbiosis in the eusocial aphid *Ceratovacuna japonica* (Hormaphidinae; Cerataphidini), involving *Buchnera* and *Arsenophonus* (24). This symbiotic partnership is consistently present across geographically distinct populations of *C. japonica* in Japan, irrespective of host plant species. The *Buchnera* genome in *C. japonica* has also undergone extreme reduction to 432,286 bp with 374 protein-coding genes, notably lacking genes for the biosynthesis of vitamin B2, riboflavin. The *Arsenophonus* genome, while larger at 853,149 bp with 532 protein-coding genes, is missing genes for synthesizing almost all essential amino acids. Nonetheless, *Arsenophonus* has retained a complete riboflavin biosynthesis pathway, suggesting metabolic complementation of the capabilities lost in *Buchnera*. Both symbionts reside within the same symbiotic organ, the bacteriome, but occupy distinct bacteriocytes. The central syncytial bacteriocytes harboring *Arsenophonus* are morphologically distinct from the peripheral uninucleate bacteriocytes harboring *Buchnera*. During embryogenesis, although a mixed population of *Buchnera* and *Arsenophonus* are transmitted maternally, they segregate into different types of bacteriocytes. This segregation indicates a high level of anatomical and developmental integration among the two symbionts and their host. Unlike Lachninae and Chaitophorinae, which often harbor *Serratia symbiotica* as a co-obligate companion symbiont, *C. japonica* has acquired *Arsenophonus*, providing an independent basis for testing whether the evolutionary dynamics of dual-symbiosis follow a common trajectory across aphid lineages. The reduced genome size of *Arsenophonus* and its anatomical and developmental integration with the host suggest a long shared evolutionary history; however, when companion symbiont acquisition and *Buchnera* genome reduction occurred within Hormaphidinae remains unclear.

In this study, we investigated the evolutionary dynamics of multi-partner symbioses in the subfamily Hormaphidinae. To comprehensively characterize the symbiotic systems in terms of symbiont diversity, genomic features, and anatomical integration, we combined 16S ribosomal RNA gene amplicon sequencing across eight Hormaphidinae species representing three tribes with detailed molecular, genomic, and microscopic analyses of two laboratory-culturable Cerataphidini species, *Ceratovacuna nekoashi* and *Pseudoregma panicola*.

## Results

### Companion symbionts coexist with *Buchnera* in *Ceratovacuna* and *Pseudoregma* aphids

To assess symbiont communities in Hormaphidinae species, we performed high-throughput 16S rRNA gene amplicon sequencing on eight species collected across Japan, representing three tribes Hormaphidini, Nipponaphidini, and Cerataphidini (Figure 1). The dominant bacterial communities in the assayed Hormaphidinae aphids comprised two α-Proteobacteria, *Hemipteriphilus* and *Rickettsia*, and four γ-Proteobacteria, *Buchnera*, *Arsenophonus*, *Gilliamella*, and *Pectobacterium* (Figure 1C). *Buchnera*, the nearly ubiquitous aphid endosymbiont, was detected in all species at extremely high relative abundance. *Arsenophonus* was found in five Cerataphidini species: *C. japonica*, *Ceratovacuna cerbera*, *C. nekoashi*, *Ceratovacuna* sp. B, and *Pseudoregma bambucicola*. *Gilliamella* was detected at low relative abundance only in *P. bambucicola*, in addition to *Buchnera* and *Arsenophonus*. *Hemipteriphilus* and *Rickettsia* were detected only from *C.* sp. B and *C. cerbera*, respectively, in addition to *Buchnera* and *Arsenophonus*, and at higher relative abundances than *Arsenophonus*. *Pectobacterium* was found only in *P. panicola*, whereas *Arsenophonus* was not detected from this species. These results demonstrate that in addition to *Buchnera*, all examined Cerataphidini species contained one or more companion symbionts with variable composition. In contrast, only *Buchnera* was detected in *Hormaphis betulae* of Hormaphidini and *Neothoracaphis yanonis* of Nipponaphidini.

**Figure 1.**
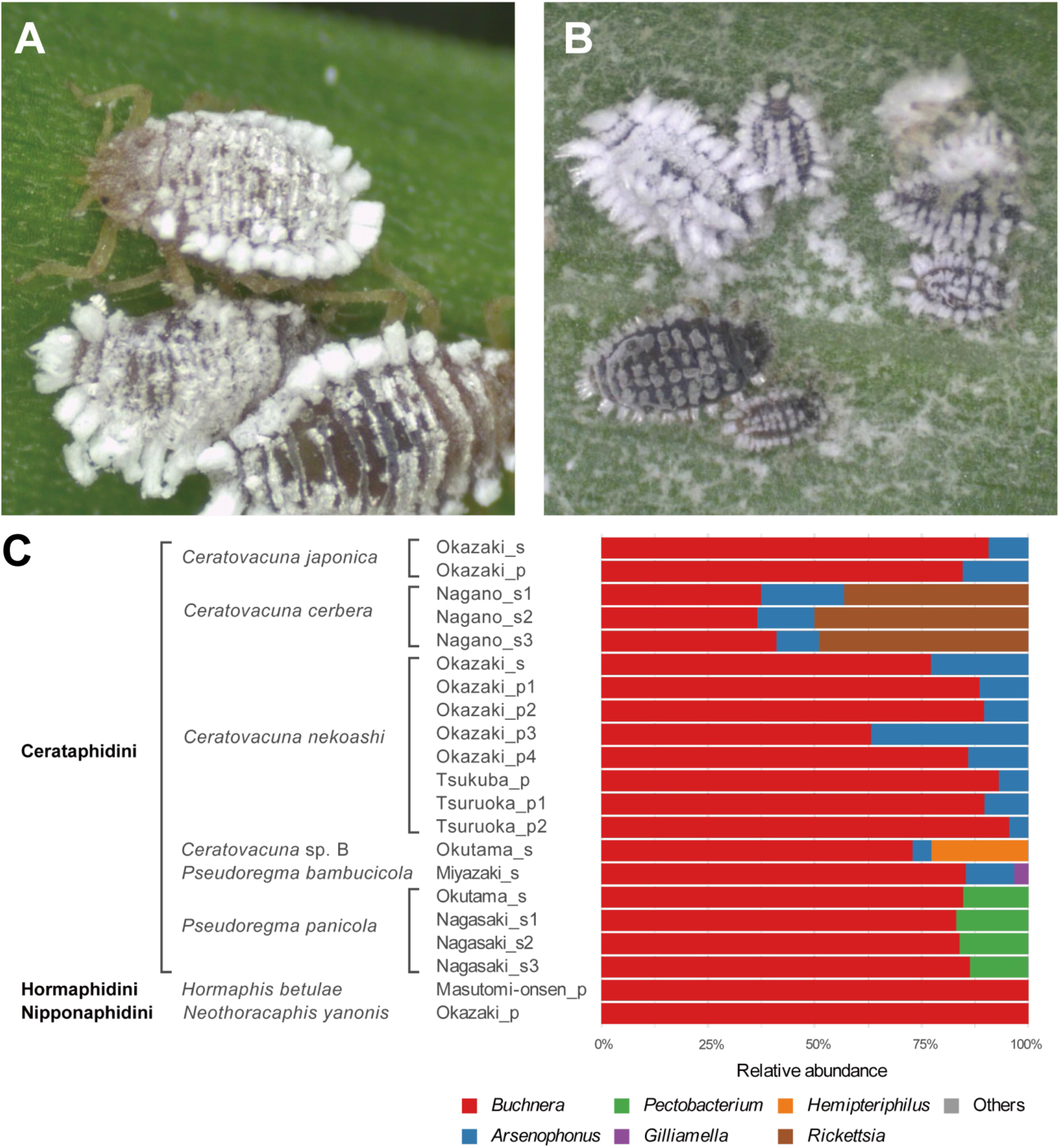
Symbiont composition in Hormaphidinae aphids revealed by 16S rRNA gene amplicon sequencing. (A) A colony of *C. nekoashi* on its secondary host plant, Japanese stiltgrass *M. vimineum*. (B) A colony of *P. panicola* on its secondary host plant, wavyleaf basketgrass *O. undulatifolius*. (C) Relative abundance of bacterial symbionts in eight Hormaphidinae species. “Others” represents sequences with an abundance of less than 1%. Labels indicate sampling locations, host plants (p: primary host, s: secondary host), and biological replicates.

Among the eight Hormaphidinae species examined, we successfully established isofemale lines of *C. nekoashi* and *P. panicola* that could be stably maintained under laboratory conditions (See details in the “Material and methods” section). These lines were cultured on the secondary host plants in the laboratory to undergo parthenogenetic reproduction, enabling detailed molecular and microscopic analyses under controlled conditions, as described below.

### Different origins of *Arsenophonus* symbionts in *C. japonica* and *C. nekoashi*

To elucidate the phylogenetic positions of bacterial symbionts in the *C. nekoashi* and *P. panicola*, we performed molecular phylogenetic analyses with our assembled data. Our phylogenetic analysis of *Buchnera*, based on 191 orthologous protein sequences, largely mirrored the relationships of their host aphid species (41, 42), with the exception of interchanged positions for the *Astegopteryx* and *Chaitoregma* genera (Figure 2A). Compared to the Aphidinae *Buchnera*, the Hormaphidini and Nippoaphidini *Buchnera* formed intermediate branches, while the six Cerataphidini *Buchnera* exhibited long branches. *Buchnera* of *C. nekoashi* was most closely related to that of *C. japonica* and formed a sister group with *Buchnera* of *P. panicola*.

**Figure 2.**
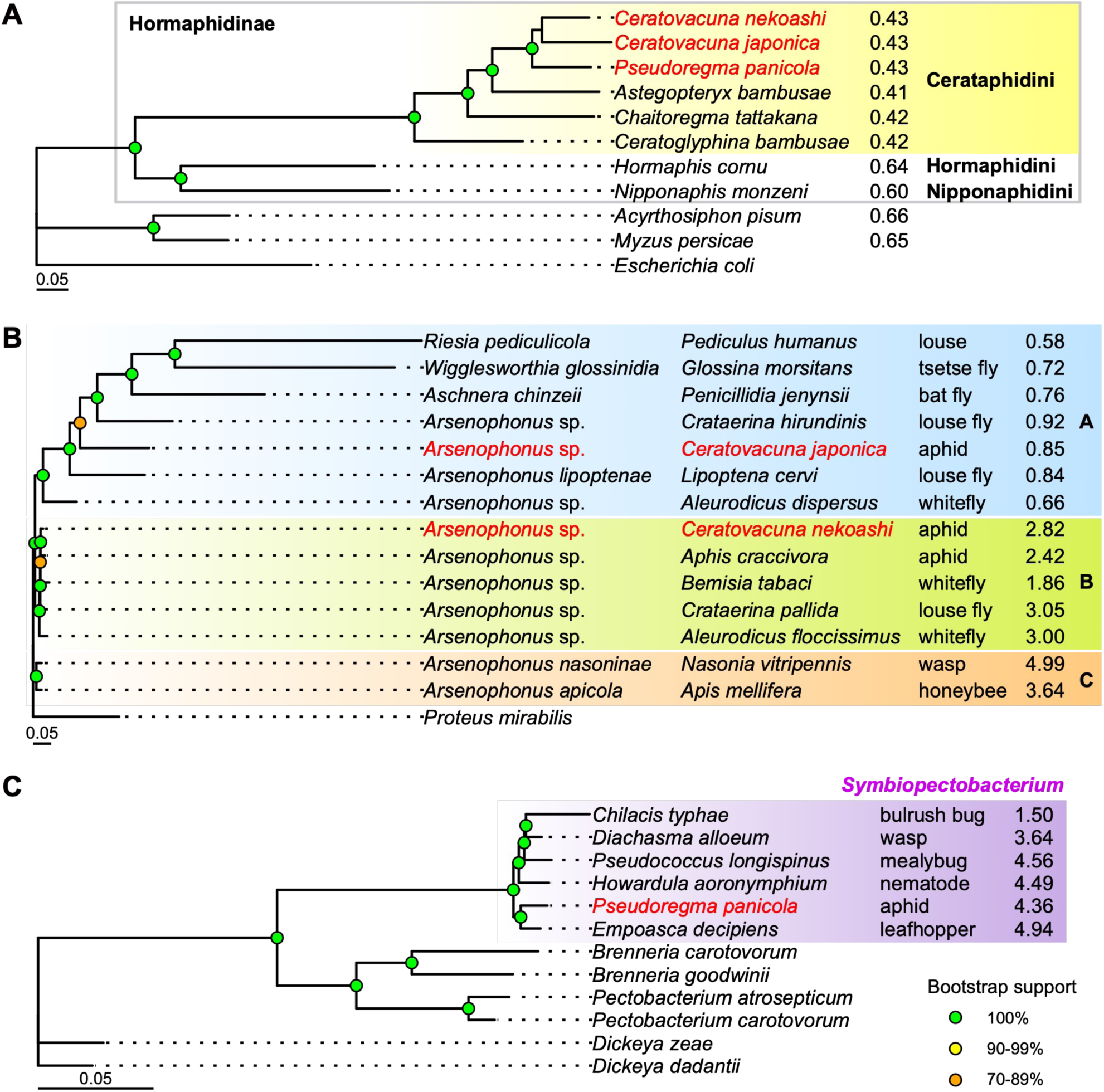
Phylogenetic analyses of *Buchnera*, *Arsenophonus*, and *Pectobacterium* (A) Maximum likelihood (ML) tree of *Buchnera aphidicola* based on 21,552 variant patterns of 42,147 amino acid positions from concatenated 191 single-copy orthologous genes. *Escherichia coli* serves as an outgroup. Labels indicate host species names. (B) ML tree of *Arsenophonus* based on 19,332 variant patterns of 41,205 amino acid positions from concatenated 131 single-copy orthologous genes. *Proteus mirabilis* serves as an outgroup. (C) ML tree of *Pectobacterium* based on 18,950 variant patterns of 150,396 amino acid positions from concatenated 483 single-copy orthologous genes. Labels in the *Symbiopectobacterium* clade indicate host species names. *Dickeya dadantii* and *Dickeya zeae* serve as outgroups. Target symbionts examined in this study are highlighted in red in each tree. Genome sizes (Mbp) of bacterial symbionts are plotted alongside taxonomic information at the tips of the trees. Bootstrap support values are indicated by colored circles at each node: green for 100%, yellow for 90-99%, and orange for 70-89%. Bootstrap values below 70% are not shown. Scale bars represent 0.05 substitutions per site.

Our phylogenetic analysis of *Arsenophonus*, based on 131 orthologous protein sequences, revealed that the *Arsenophonus* symbionts in *C. japonica* and *C. nekoashi* belong to different phylogenetic clades (Figure 2B). The analysis identified three clades: clade A primarily contained symbionts from a louse (Psocodea) and hematophagous Diptera, including *Arsenophonus* and related genera such as *Riesia*, *Wigglesworthia*, and *Aschnera*, all characterized by intermediate to long branch lengths and reduced genomes smaller than 1 Mbp; clade B predominantly included *Arsenophonus* symbionts from Hemiptera with intermediate genome size ranging from 1.86 to 3.05 Mbp; and clade C exclusively comprised *Arsenophonus* from Hymenoptera with large genomes ranging from 3.64 to 4.99 Mbp. The *Arsenophonus* of *C. japonica* belonged to clade A, while that of *C. nekoashi* belonged to clade B. This phylogenetic difference contrasts with the *Buchnera* phylogeny, where *C. japonica* and *C. nekoashi* grouped within the tribe Cerataphidini (Figure 2A).

We constructed an additional phylogenetic tree of the *Arsenophonus* symbionts with 16S rRNA gene sequences, including those of *C. cerbera*, *C.* sp. B, *P. bambucicola*, and two additional Cerataphidini species, *Cerataphis bambusifoliae* and *Astegopteryx bambusae*, reported in Xu et al. (43). This analysis revealed that the *Arsenophonus* of *C. japonica* fell within clade A, whereas those of the remaining six Cerataphidini species fell within clade B (Figure S1A). A multiple sequence alignment further showed that the clade B *Arsenophonus* sequences were relatively conserved across the six Cerataphidini species, while that of *C. japonica* exhibited numerous variant sites (Figure S1B).

### Pectobacterium symbiont of P. panicola belongs to the Symbiopectobacterium clade

Our phylogenetic analysis of *Pectobacterium* and related bacteria, based on 483 orthologous protein sequences, revealed that the *Pectobacterium* symbiont of *P. panicola* is phylogenetically positioned within the *Symbiopectobacterium* clade (Figure 2C). The *Pectobacterium* symbiont of *P. panicola* is the most closely related to *Symbiopectobacterium purcellii*, previously isolated from the leafhopper *Empoasca decipiens* (44). This phylogenetic placement supports the reclassification of this symbiont as a member of the *Symbiopectobacterium* genus. Consequently, we propose to refer to this bacterial symbiont as *Symbiopectobacterium* sp.

### Intracellular localization of *Buchnera* and *Arsenophonus* in *C. nekoashi*

We characterized the spatial organization of bacterial symbionts in *C. nekoashi*, a congener of *C. japonica*. Fluorescence *in situ* hybridization (FISH) analysis of cryosections from parthenogenetic viviparous third/fourth instar nymphs revealed paired symbiotic organs (bacteriomes) in the abdomen (Figure 3A). The bacteriomes in the anterior abdomen exhibited an oval shape, while the bacteriomes in the posterior abdomen were longitudinal and laterally positioned. In both bacteriome types, *Buchnera* was localized within large rounded bacteriocytes, while *Arsenophonus* was localized within sheath cells interspersed among the *Buchnera*-containing bacteriocytes (Figure 3B, 3C). No extracellular signals were detected for either symbiont.

**Figure 3.**
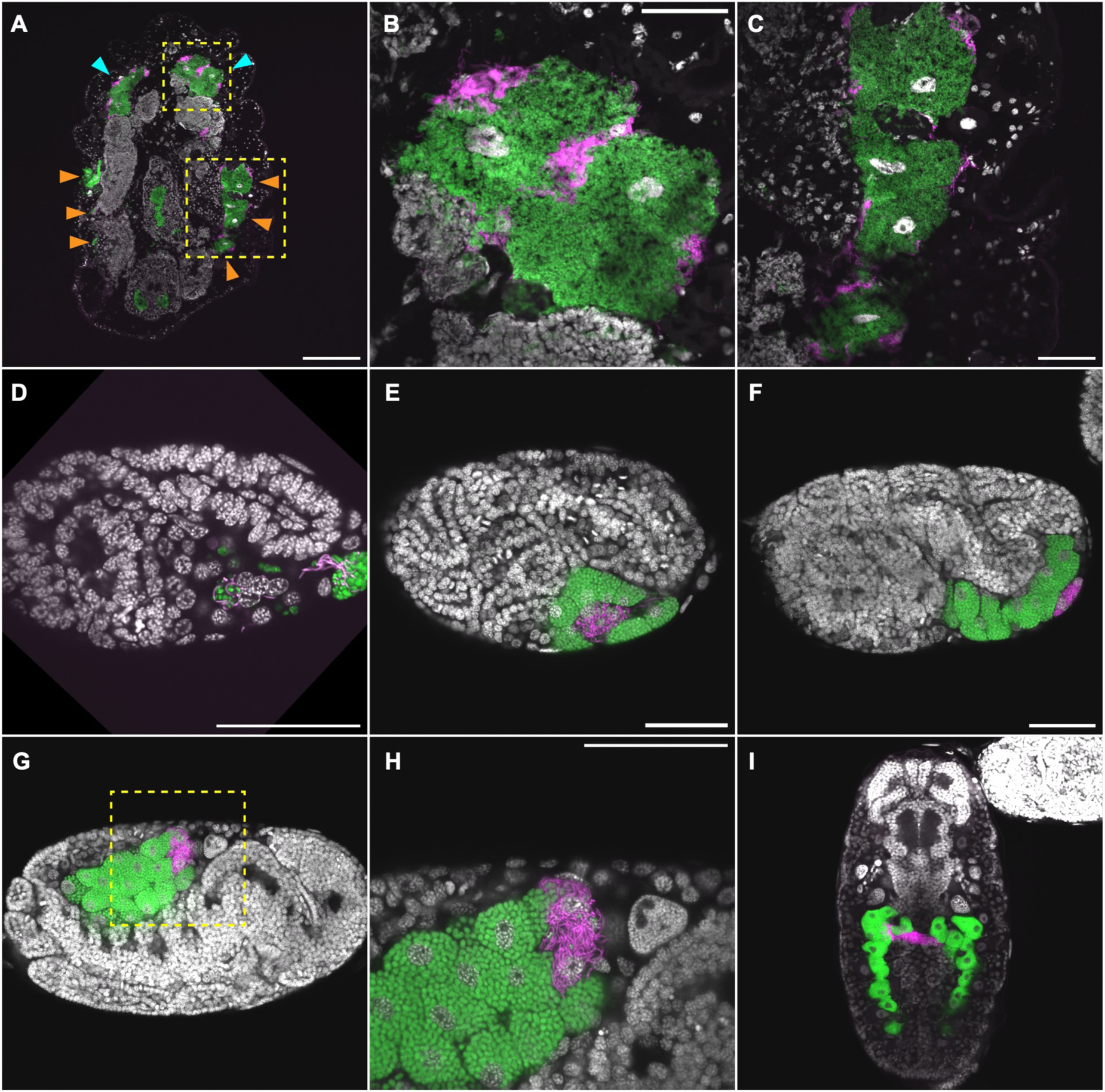
Localization of *Buchnera* and *Arsenophonus* symbionts in *C. nekoashi* (A–C) FISH images of cryosections from a third or fourth instar nymph. (A) Whole body. Cyan arrowheads indicate anterior bacteriomes and orange arrowheads indicate posterior bacteriomes. Yellow dotted rectangles indicate regions magnified in anterior bacteriomes (B) and posterior bacteriomes (C). (D–I) FISH images of bacteriome formation during embryogenesis. (D) Stage 12 embryo. Both *Buchnera* and *Arsenophonus* are incorporated into the posterior pole of the embryo. (E) Stage 13 embryo. Peripheral uninucleate bacteriocytes harboring *Buchnera* are formed, while *Arsenophonus* co-resides with *Buchnera* in a central syncytial bacteriocyte. (F) Stage 14 embryo. During germ band extension, *Arsenophonus* occupies a syncytial bacteriocyte at the periphery of the bacteriome. (G) Stage 17 embryo. The bacteriome is repositioned to the dorsal abdomen after katatrepsis. (H) Higher magnification of the boxed region in (G). A syncytial bacteriocyte contains predominantly filamentous *Arsenophonus* with a small number of *Buchnera* cells, while uninucleate bacteriocytes harbor exclusively spherical *Buchnera*. (I) Stage 19–20 embryo. The bacteriome exhibits a horseshoe-shaped structure, with the *Arsenophonus*-containing syncytial bacteriocyte extending laterally. Nuclei (DAPI) in gray; *Buchnera* in green (Cy5); *Arsenophonus* in magenta (Cy3). Scale bars: 200 µm (A), 50 µm (B–I).

We next examined the symbiont localization during embryonic development, following the developmental staging system established for the pea aphid by Miura et al. (16). In stage 12 embryos, characterized by progressive segmentation, both *Buchnera* and *Arsenophonus* were incorporated into the posterior pole (Figure 3D). By stage 13, coinciding with limb bud formation, bacteriocytes had begun to form ventrally: *Buchnera* was localized within peripheral uninucleate bacteriocytes, while *Arsenophonus* co-resided with *Bcuhnera* in a central syncytial bacteriocyte (Figure 3E). At stage 14, during germ band extension, *Arsenophonus* occupied a syncytial bacteriocyte at the periphery of the bacteriome (Figure 3F). After katatrepsis, by stage 17, the bacteriome was repositioned to the dorsal abdomen (Figure 3G). At this stage, spherical *Buchnera* occupied uninucleate bacteriocytes, while filamentous *Arsenophonus* resided in a syncytial bacteriocyte also containing a small number of *Buchnera* cells (Figure 3I). By stage 19–20, prior to larviposition, the bacteriome exhibited a horseshoe-shaped structure, with the *Arsenophonus*-containing syncytial bacteriocyte extending laterally as a bridge-like structure (Figure 3I). The presence of both symbionts in embryos demonstrates their vertical maternal transmission.

### Intracellular localization of Buchnera and Symbiopectobacterium in P. panicola

We next examined *P. panicola*, which harbors *Symbiopectobacterium* as a companion symbiont. FISH analysis of cryosections from parthenogenetic viviparous third/fourth instar nymphs revealed paired elongated bacteriomes in the anterior abdomen (Figure 4A). Within these bacteriomes, *Buchnera* was localized within large bacteriocytes, while *Symbiopectobacterium* was localized within bacteriocytes embedded among the *Buchnera*-containing bacteriocytes (Figure 4C). No extracellular signals were detected for either symbiont.

**Figure 4.**
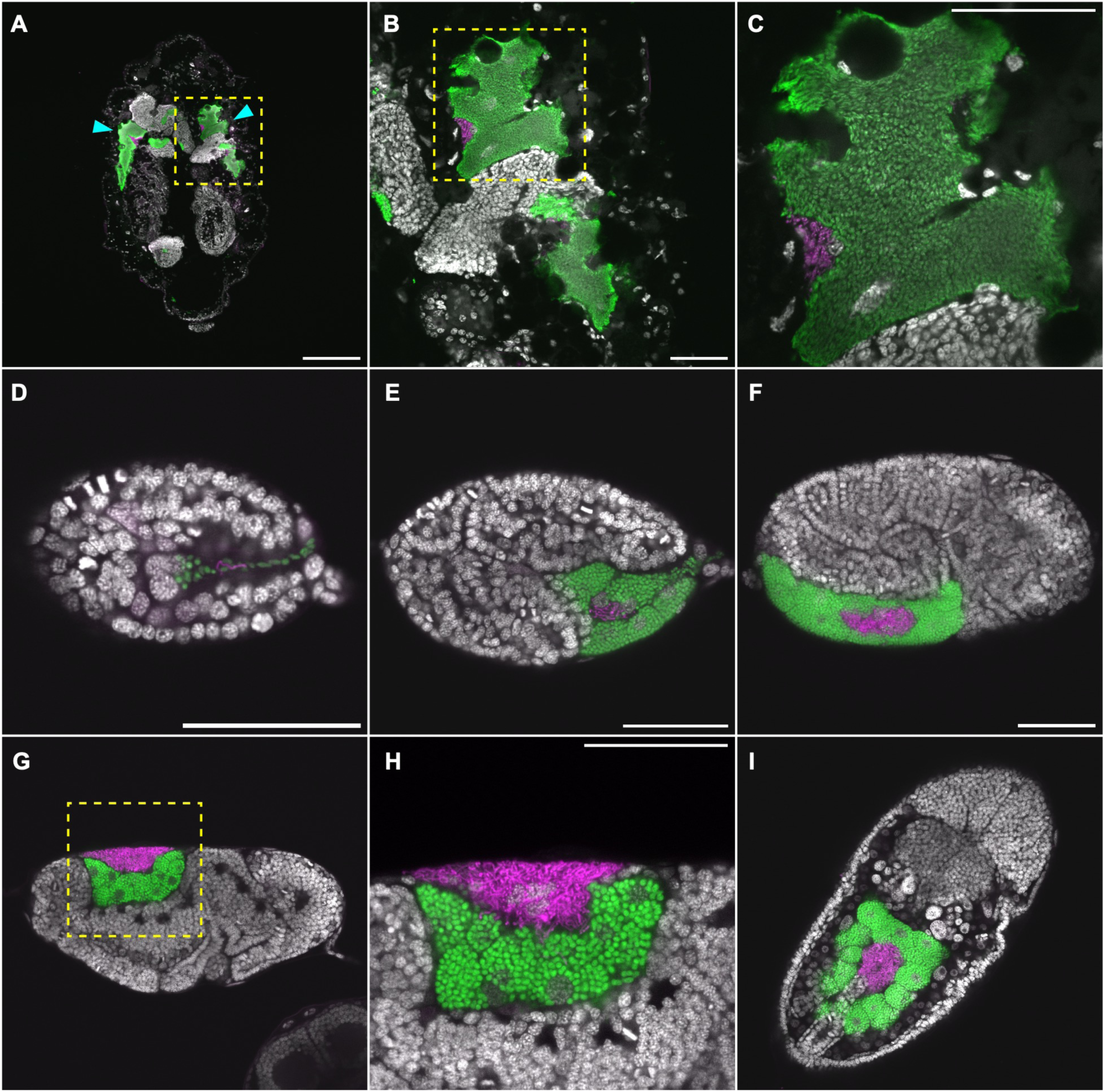
Localization of *Buchnera* and *Symbiopectobacterium* symbionts in *P.* panicola. (A–C): FISH images of cryosections from a third or fourth instar nymph. (A) Whole body. Cyan arrowheads indicate anterior bacteriomes. (B) Higher magnification of the boxed region in (A). (C) Higher magnification of the boxed region in (B). (D–I) FISH images of bacteriome formation during embryogenesis. (D) Stage 8 embryo. Both *Buchnera* and *Symbiopectobacterium* are incorporated together into the posterior pole. (E) Stage 13 embryo. Peripheral uninucleate bacteriocytes harboring *Buchnera* are formed, while *Symbiopectobacterium* co-resides with *Buchnera* in a central syncytial bacteriocyte. (F) Stage 14 embryo. Symbiopectobacterium is localized within the central syncytial bacteriocyte, while Buchnera occupies the peripheral uninucleate bacteriocytes. (G) Lateral view of Stage 17 embryo. The bacteriome is repositioned to the dorsal abdomen after katatrepsis. (H) Higher magnification of the boxed region in (G). Uninucleate bacteriocytes harbor spherical *Buchnera* cells, while a large syncytial bacteriocyte at the periphery contains filamentous *Symbiopectobacteirum* cells. (I) Dorsal view of Stage 17 embryo. The *Symbiopectobacterium*-containing syncytial bacteriocyte is centrally positioned within the bacteriome. Nuclei (DAPI) in gray; *Buchnera* in green (Cy5); *Symbiopectobarium* in magenta (Cy3). Scale bars: 200 µm (A), 50 µm (B–I).

We next examined symbiont localization during embryonic development. In stage 8 embryos, during anatrepsis, both *Buchnera* and *Symbiopectobacterium* were incorporated together into the embryo from the posterior pole (Figure 4D). By stage 13, coinciding with limb bud formation, peripheral uninucleate bacteriocytes harboring *Buchnera* had formed ventrally, while *Symbiopectobacterium* co-resided with *Buchnera* in a central syncytial bacteriocyte (Figure 4E). At stage 14, during germ band extension, *Symbiopectobacterium* was localized within the central syncytial bacteriocyte, while *Buchnera* occupied the peripheral uninucleate bacteriocytes (Figure 4F). After katatrepsis, by stage 17, the bacteriome was repositioned to the dorsal abdomen (Figure 4G). In lateral view, spherical *Buchnera* cells occupied uninucleate bacteriocytes, while filamentous Symbiopectobacterium cells resided in a large syncytial bacteriocyte at the periphery of the bacteriome (Figure 4G, 4H). In dorsal view, the *Symbiopectobacterium*-containing syncytial bacteriocyte was centrally positioned within the bacteriome (Figure 4I). The presence of both symbionts throughout embryogenesis demonstrates vertical maternal transmission.

### Streamlined *Buchnera* genomes with consistent riboflavin complementarity patterns in C. nekoashi and P. panicola

We determined the complete genome sequences of *Buchnera* from both *C. nekoashi* and *P. panicola* and the *Arsenophonus* from *C. nekoashi*, as well as the nearly complete genome sequences of *Symbiopectobacterium* from *P. panicola*. Both *Buchnera* genomes comprise one circular chromosome and two plasmid, pLeu and pTrp (Table 1, Figure 5A, 6A). The *Buchnera* chromosomes of *C. nekoashi* and *P. panicola* are comparable in size (416,450 bp and 412,843 bp) and GC content (19.5% and 19.0%), and are also similar in size to that of the closely related species *C. japonica* (414,725 bp) (24). These genome sizes are significantly smaller than ∼0.6 Mb *Buchnera* genomes in mono-symbioses and fall within the ∼0.4 Mb *Buchnera* genomes in co-obligate symbioses (10, 13, 19–22, 24, 45). Both *Buchnera* genomes encode similar number of protein-coding genes (361 and 363 CDSs), with identical numbers of rRNAs (3) and tRNAs (30). The plasmid pLeu and pTrp showed identical gene arrangements in both species (*leuBCDA*, *yghA*, and *repA* genes in pLeu and one *trpE*, one pseudogenized *trpG*, and 10 *trpG* genes in pTrp), mirroring those of *C. japonica* (24). Genome-wide synteny analysis revealed highly conserved gene order among *Buchnera* chromosomes of *A. pisum*, *C. japonica*, *C. nekoashi*, and *P. panicola* (Figure S2A). Notably, the three *Buchnera* genomes of *C. japonica*, *C. nekoashi*, and *P. panicola* commonly lost 201 orthologous genes compared to that of *A. pisum* (Table S3), including genes involved in ornithine synthesis (*argA*–*E*), riboflavin synthesis (*ribA*, *ribB*, *ribD*, *ribE*, *ribH*, *yigL*), peptidoglycan synthesis (*glmM*, *glmS*, *glmU*, *murA*–*G*, *mraY*, *mrcB*, *ftsI*), flagellar components (*flgA*, *flgD*, *flgK*, *fliK*, *fliM*), chromosomal replication (*dnaA*), and DNA mismatch repair (*mutL*, *mutS*).

**Figure 5.**
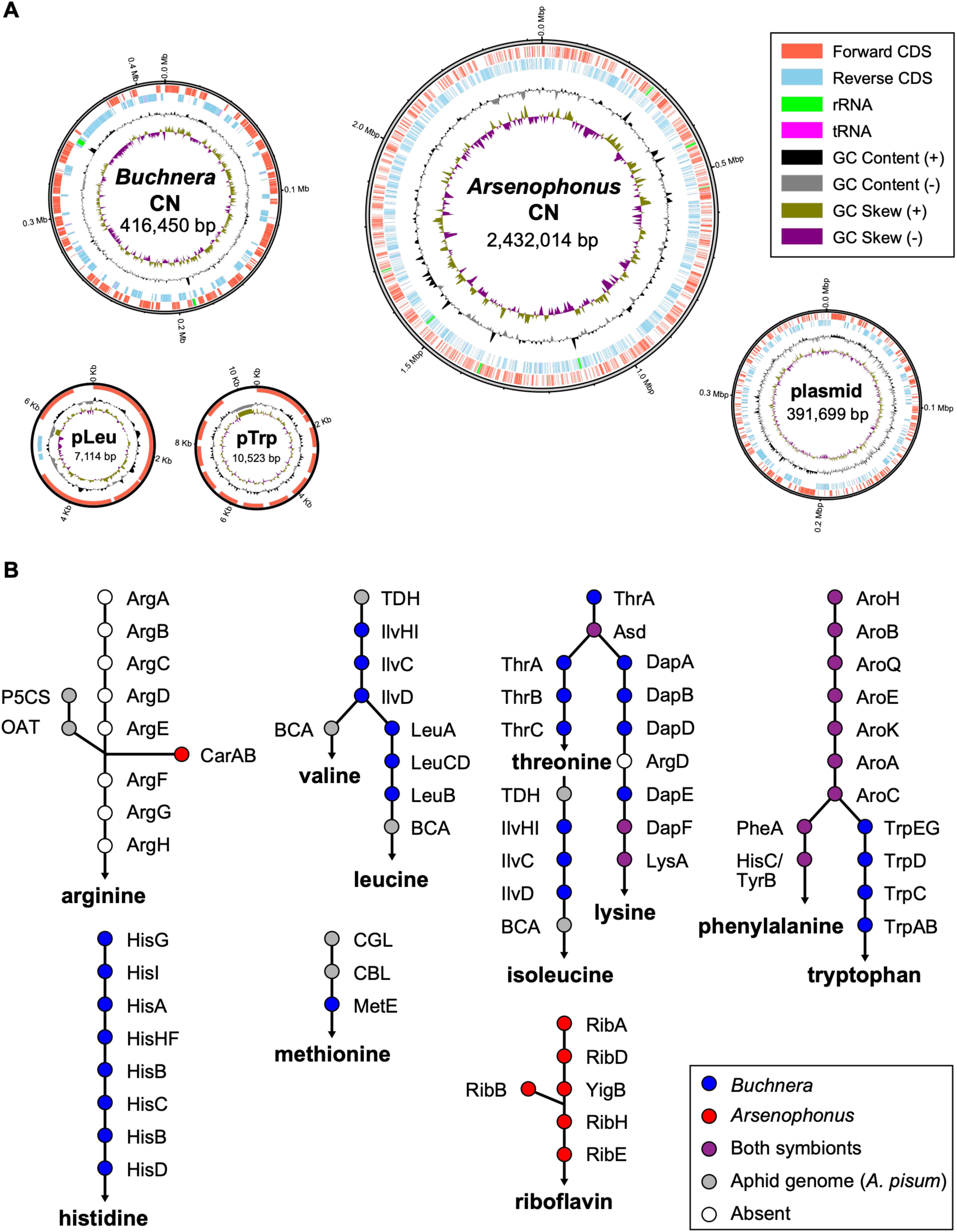
Genomic features and metabolic gene repertoires of symbionts in *C.*nekoashi. (A) Circular genome maps of *Buchnera* and *Arsenophonus* of *C. nekoashi*. Tracks from outside to inside: forward CDS (orange), reverse CDS (cyan), rRNA (green), tRNA (magenta), GC content, and GC skew. Genome sizes are indicated at the center. (B) Gene repertoires for essential amino acid and riboflavin biosynthesis pathways in *C. nekoashi*. Filled circles indicate genes present in *Buchnera* (blue), *Arsenophonus* (red), or both symbionts (purple). Gray circles indicate genes presumably encoded in the host aphid genome, based on the *A. pisum* genome. Open circles indicate absent genes.

**Table 1.** General features of sequenced symbiont genomes of *C. nekoashi*, *P. panicola*, and the closely related species. *B. aphidicola* of *C. japonica* (GCA_024349705.1); *B. aphidicola* of *A. pisum* (GCA_000009605.1); *Arsenophonus* sp. of *C. japonica* (GCA_024349725.1); *S. purcellii* of *E. decipiens* (GCA_019797845.1). These genomes were re-annotated using DFAST with identical parameters to ensure consistent comparison with our target symbionts.

| Symbiont | Host | Genome size (bp) | Chromosome size (bp) | Number of contigs | Plasmid | GC (%) | Coding density (%) | Intact CDSs | tRNA | rRNA | Pseudo-genes | Assembly level |
| --- | --- | --- | --- | --- | --- | --- | --- | --- | --- | --- | --- | --- |
| *Buchnera* | *C. japonica* | 432,286 | 414,725 | 3 | 2 (pLeu, pTrp) | 20.0 | 88.1 | 374 | 30 | 3 | 29 | complete |
|  | *C. nekoashi* | 434,087 | 416,450 | 3 | 2 (pLeu, pTrp) | 19.5 | 85.6 | 361 | 30 | 3 | 42 | complete |
|  | *P. panicola* | 430,341 | 412,843 | 3 | 2 (pLeu, pTrp) | 19.0 | 88.7 | 363 | 30 | 3 | 45 | complete |
|  | *A. pisum* | 655,725 | 640,681 | 3 | 2 (pLeu, pTrp) | 26.3 | 87.2 | 571 | 31 | 3 | 24 | complete |
| *Arsenophonus* | *C. japonica* | 853,149 | 853,149 | 1 | 0 | 18.5 | 60.6 | 532 | 32 | 3 | 23 | complete |
|  | *C. nekoashi* | 2,823,713 | 2,432,014 | 2 | 1 | 37.0 | 73.2 | 2,340 | 60 | 22 | 1,094 | complete |
| *Symbiopectobacterium* | *P. panicola* | 4,358,940 | 4,358,940 | 2 | 0 | 52.9 | 83.7 | 2,963 | 57 | 19 | 1,993 | contigs |
|  | *E. decipiens* | 4,942,431 | 4,942,431 | 1 | 0 | 52.5 | 85.5 | 4,061 | 73 | 21 | 609 | complete |

The *Arsenophonus* genome of *C. nekoashi* consisted of one circular chromosome (2,432,014 bp, 37.0% GC content) and one large circular plasmid (Table 1, Figure 5A). This genome size is similar to that of the facultative *Arsenophonus* symbiont in *Aphis craccivora* (2,424,437 bp, ASM1346013v1) and significantly larger than that of the co-obligate *Arsenophonus* symbiont in *C. japonica* (853,149 bp) (24). The *Arsenophonus* genome contained numerous pseudogenes (1,094) and exhibited lower coding density (73.2%) than a typical 85–90% range for bacterial genomes (46), indicating ongoing genomic erosion. Genome-wide synteny analysis revealed extensive genome rearrangements between the *Arsenophonus* chromosomes of *C. japonica* and *C. nekoashi* (Figure 4B), consistent with their placement in different phylogenetic clades (Figure 2B). Substantial rearrangements were also observed between *C. nekoashi* and *A. craccivora*, despite their similar genome sizes and close phylogenetic placement within clade B (Figure S2B).

The *Symbiopectobacterium* genome of *P. panicola* was assembled into two contigs totaling 4,358,940 bp with 52.9% GC content (Table 1, Figure 6A), representing the largest genome among all known secondary symbionts in aphids (47). Although a complete circular genome could not be obtained due to low read depth (∼8x) and long repeats, the genome size is comparable to that of *Symbiopectobacterium purcellii* from the leafhopper *Empoasca decipiens* (4,942,431 bp) (44). The *Symbiopectobacterium* genome contained numerous pseudogenes (1,993) and exhibited relatively low coding density (83.7%), indicating ongoing genomic erosion.

**Figure 6.**
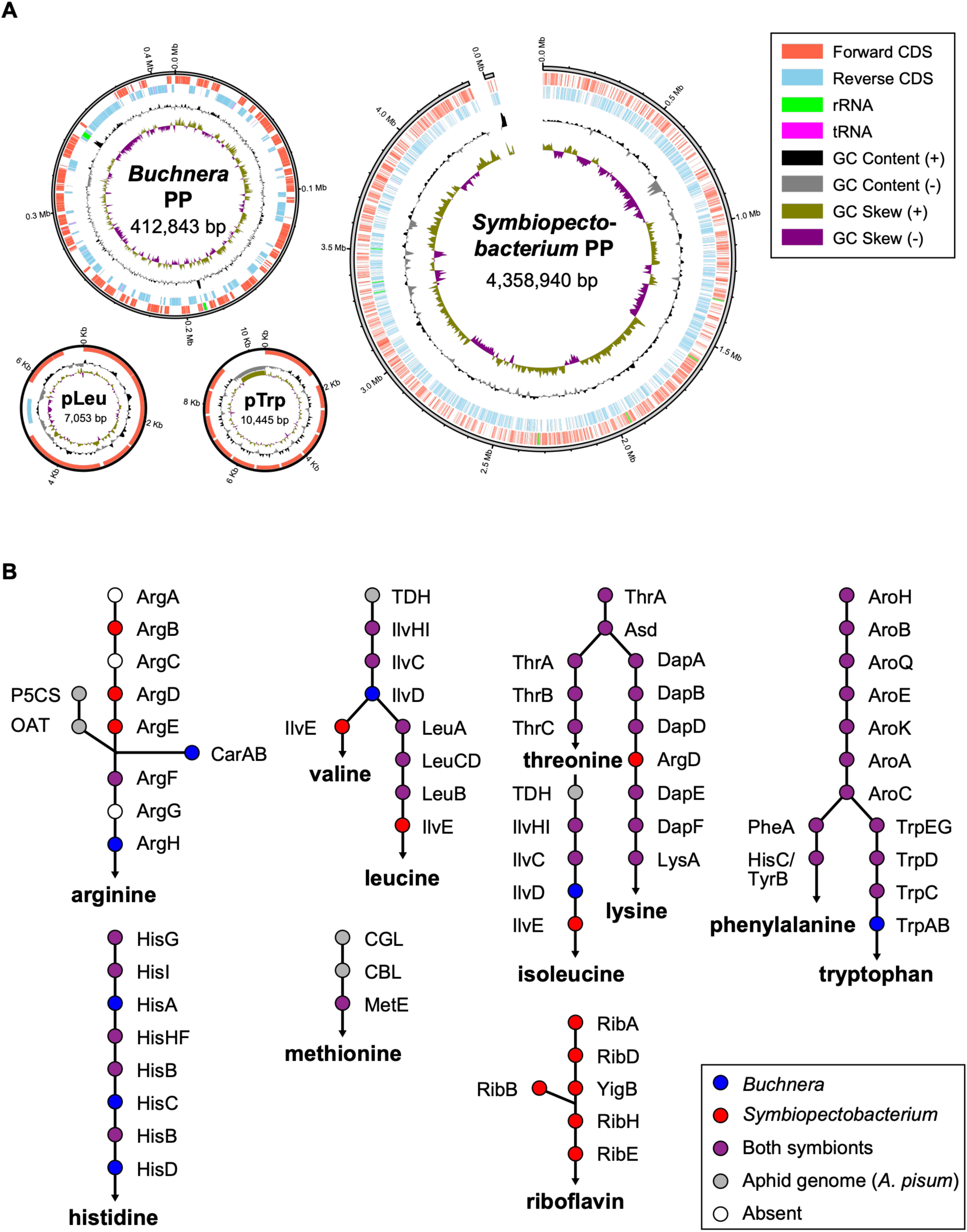
Genomic features and metabolic gene repertoires of symbionts in *P.* panicola. (A) Circular genome maps of *Buchnera* and *Symbiopectobacterium* of *P. panicola*. Tracks from outside to inside: forward CDS (orange), reverse CDS (cyan), rRNA (green), tRNA (magenta), GC content, and GC skew. Genome sizes are indicated at the center. (B) Gene repertoires for essential amino acid and riboflavin biosynthesis pathways in *P. panicola*. Filled circles indicate genes present in *Buchnera* (blue), *Symbiopectobacterium* (red), or both symbionts (purple). Gray circles indicate genes presumably encoded in the host aphid genome, based on the *A. pisum* genome. Open circles indicate absent genes.

In aphids with dual symbiotic systems, the ∼0.4 Mb *Buchnera* genomes have frequently lost genes involved in various biosynthesis pathways, while companion symbionts often retain these genes, suggesting potential metabolic complementation (19–24). In both *C. nekoashi* and *P. panicola*, the *Buchnera* genomes lack all genes responsible for riboflavin biosynthesis, while these genes are present in the *Arsenophonus* and *Symbiopectobacterium* genomes (Figure 5B, 6B). This suggests that the companion symbionts have the genetic capacity to complement this biosynthetic deficiency in *Buchnera*. This pattern is consistent with the *Buchnera*–*Arsenophonus* dual symbiosis observed in *C. japonica* (24). On the other hand, complementarity is incomplete for arginine biosynthesis. In *C. japonica*, the *Buchnera* genome retains *argF*, *argG*, and *argH* genes, which can convert ornithine to arginine. In contrast, the *Buchnera* of *C. nekoashi* has lost all arginine biosynthesis genes, and its *Arsenophonus* only contains a pseudogenized *argE*. In *P. panicola*, *Buchnera* retains intact *argF* and *argH*, but *argG* is pseudogenized. While *Symbiopectobacterium* retains intact *argF,* its *argG* is also pseudogeneized and *argH* is absent. As *argG* is non-functional in both symbionts, the pathway for converting ornithine to arginine remains incomplete.

## Discussion

### *Buchnera* genome reduction into ∼0.4 Mbp in the tribe Cerataphidini

The genome sizes of *Buchnera* in Cerataphidini aphids, including *C. japonica*, *C. nekoashi*, and *P. panicola* are significantly small at approximately 0.4 Mbp (Table 1). Recent studies have revealed that *Buchnera* in several other Cerataphidini genera, including *Astegopteryx bambusae*, *Chaitoregma tattakana*, and *Ceratoglyphina bambusae*, also possess extremely reduced genomes of ∼0.4 Mbp (48). In contrast, *Buchnera* genomes in other Hormaphidinae tribes retain larger sizes typical of mono-symbiotic systems, such as 0.64 Mbp in *Hormaphis cornu* from Hormaphidini and 0.60 Mbp in *Nipponaphis monzeni* from Nipponaphidini (49, 50). The highly conserved gene order across *Buchnera* chromosomes (Figure S2A) and identical gene arrangements in pLeu and pTrp plasmids suggest coordinated genome reduction events. Moreover, the identical pattern of gene losses among *Buchnera* in *C. japonica*, *C. nekoashi*, and *P. panicola* (Figure 5B, 6B, and Table S3), including genes involved in various metabolic pathways and cellular processes, indicates that these losses occurred before the divergence of these lineages. Collectively, these findings suggest that the drastic genome reduction into ∼0.4 Mbp occurred in the common ancestor of Cerataphidini *Buchnera*.

Despite their reduced genome size of ∼0.4 Mbp, the *Buchnera* in *C. nekoashi* and *P. panicola* have retained genes responsible for synthesizing most essential amino acids (Figure 5B, 6B), suggesting their maintained role in host nutrition. In contrast, these *Buchnera* have lost most genes for riboflavin and arginine biosynthesis pathways. While companion symbionts in both aphids possess complete riboflavin biosynthesis pathways, providing the genetic capacity for this biosynthesis, arginine biosynthesis remains incomplete. Such incomplete complementarity patterns have been also observed in other aphid dual-symbioses (10, 22). Our understanding of aphid nutrient requirements has primarily been based on the studies of *A. pisum*, involving analyses of phloem sap nutrients, *Buchnera* elimination experiments, and comparative genomics between *Buchnera* and the host (13, 51–54). Meanwhile, nutritional requirements can vary among aphid biotypes or species (55, 56). Similarly, *C. nekoashi* and *P. panicola* may have evolved nutritional requirements that differ from those of *A. pisum*, but further studies are needed to confirm this.

### Roles of the companion symbionts in the tribe Cerataphidini

Genomic analyses revealed that companion symbionts likely complement the highly reduced *Buchnera* genomes in Cerataphidini. The *Arsenophonus* in *C. nekoashi* appears to function primarily as a riboflavin provider based on its complete biosynthesis pathways (Figure 5B), which is consistent with *Arsenophonus* functions reported in other Hemiptera and Diptera (24, 57–59). The *Symbiopectobacterium* in *P. panicola* also appears to function primarily as a riboflavin provider based on its complete biosynthesis pathway (Figure 6B). Additionally, it has retained complete pathways for several essential amino acids, leucine, lysine, phenylalanine, and threonine, that overlap with *Buchnera* functions, potentially providing a redundant metabolic capacity to buffer against functional losses resulting from *Buchnera*’s accelerated evolutionary rate (60). Notably, our genomic analyses revealed that *Buchnera* in *C. nekoashi* and *P. panicola* have lost crucial genes for DNA replication and repair, including *dnaA*, *mutL*, and *mutS*, while these genes are retained in their companion symbionts. A recent study has demonstrated that a facultative symbiont *Serratia* can support *Buchnera* genome stability through its DNA mismatch repair system in *A. pisum* (61). This suggests that companion symbionts may similarly contribute to maintaining *Buchnera* functionality, indirectly ensuring the survival and fitness of the host aphids.

Most Cerataphidini species alternate between primary hosts on *Styrax* trees and secondary hosts across several unrelated plant families (62). For instance, *C. japonica* and *C. nekoashi* share the same primary host *Styrax japonica*, but they differ in their secondary hosts: *C. japonica* colonizes bamboo grasses in the Bambusoideae such as *Sasa senanensis*, *Pleioblastus chino*, and *Pleioblastus simonii*, whereas *C. nekoashi* colonizes the Panicoideae grass *M. vimineum*. Such host plant switching necessitates mechanisms to cope with diverse plant defense responses, including secondary metabolites (63). Notably, these two Cerataphidini species harbor phylogenetically distinct *Arsenophonus* strains, suggesting a potential role of symbionts in host adaptation. A previous study demonstrated that *Arsenophonus* in the polyphagous aphid *A. craccivora* is associated with an expanded host plant range (64). In addition, *Arsenophonus* clusters with phloem-restricted plant pathogens such as *Candidatus* Phlomobacter fragariae and *Candidatus* Arsenophonus phytopathogenicus (65). Collectively, these findings suggest that the companion symbionts might contribute to counteracting plant defenses and enhancing host plant utilization capabilities in Cerataphidini aphids.

### Organization of the bacteriome morphology differs in dual-symbiotic systems

The localization of companion symbionts in dual-symbiosis exhibits remarkable diversity, reflecting the complex evolutionary trajectories of these symbiotic relationships. In *C. japonica*, *Arsenophonus* resides inside the central, large syncytial bacteriocytes through all developmental stages (24). In *C. nekoashi*, *Arsenophonus* similarly occupies a syncytial bacteriocyte during embryogenesis, but localizes in sheath cells in the nymphal stage (Figure 3). This developmental stage-specific relocalization is reminiscent of facultative *Serratia symbiotica* in *A. pisum*, where *Serratia* inhabits secondary bacteriocytes during embryogenesis but also localizes in sheath cells at later stages (66). In *P. panicola*, *Symbiopectobacterium* resides in a syncytial bacteriocyte during embryogenesis and remains in a single bacteriocyte within each bacteriome in the nymphal stage (Figure 4). These diverse patterns suggest ongoing integration processes for recently acquired symbionts into host developmental pathways. Comparable diversity has been documented in other aphid lineages, such as Lachninae and Chaitophorinae aphids (67, 68). For instance, the co-obligate *Serratia symbiotica* in *Periphyllus lyropictus* exhibits both intracellular and extracellular localization, including in hemolymph and gut (68). This localization diversity reflects the dynamic nature of host-symbiont interactions and suggests a potential evolutionary continuum from facultative to obligate symbiosis.

### Frequent replacements of companion symbionts across dual-symbiotic systems

All examined *Ceratovacuna* species and *P. bambucicola* harbored *Arsenophonus* alongside the primary endosymbiont *Buchnera* (Figure 1C). Our phylogenetic analysis revealed that clade B *Arsenophonus* are closely related across diverse Cerataphidini species, including *C. cerbera*, *C.* sp. B, *C. nekoashi*, *P. bambucicola*, *C. bambusifoliae*, and *A. bambusae*, and are also closely related to *Arsenophonus* found in non-Cerataphidini insects (Figure S1). In contrast, *C. japonica* harbors a phylogenetically distinct clade A *Arsenophonus* with a more reduced genome (853,149 bp) compared to the clade B *Arsenophonus* of *C. nekoashi* (2,432,014 bp) (Figure 2B). These findings suggest that the clade A *Arsenophonus* in *C. japonica* represents an ancestral symbiont, while clade B *Arsenophonus* strains were independently acquired in other Cerataphidini lineages (Figure 7).

**Figure 7.**
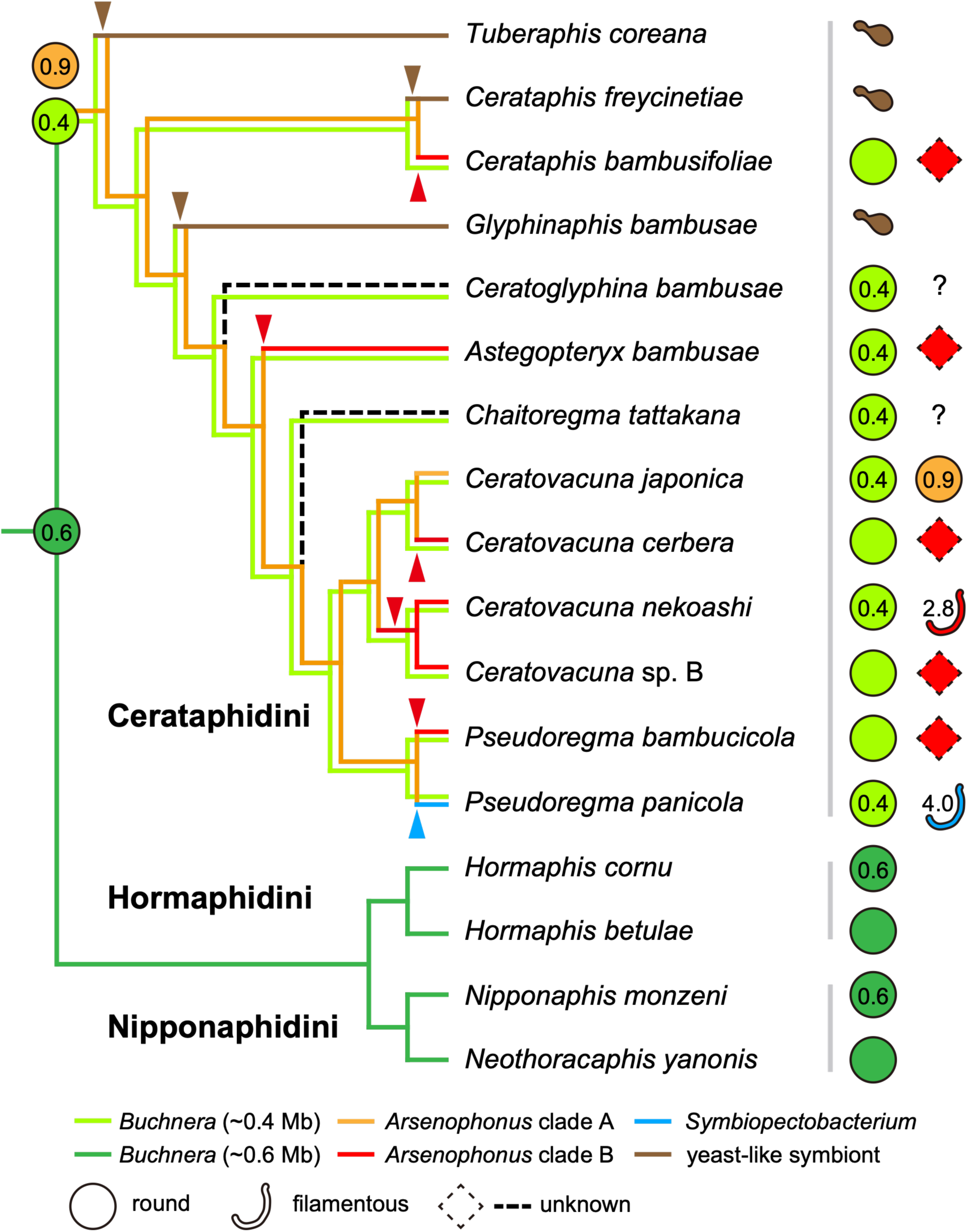
Proposed evolutionary trajectory of multi-partner endosymbiosis in the subfamily Hormaphidinae. Cladogram depicts the phylogenetic relationship among Hormaphidinae species based on (41, 42). Symbiont associations are represented by colored branches: *Buchnera* with ∼0.4 Mb genome (light green); *Buchnera* with ∼0.6 Mb genome (dark green); *Arsenophonus* clade A (orange); *Arsenophonus* clade B (red); and *Symbiopectobacterium* (cyan). Color transitions along branches indicate presumed evolutionary events: extreme genome reduction to ∼0.4 Mb in *Buchnera* (dark to light green), replacement of the ancestral clade A *Arsenophonus* by clade B *Arsenophonus* (orange to red) in most Cerataphidini lineages, and replacement by *Symbiopectobacterium* (orange to cyan) in *P. panicola*.

In *P. panicola*, we found *Symbiopectobacterium* instead of *Arsenophonus* in addition to *Buchnera* (Figure 1C). Notably, previous studies have reported different companion symbionts in *P. panicola* from other geographical populations: *Wolbachia* in a Taiwanese population and *Gilliamella* in a Thai population (10). Furthermore, Chinese *P. bambucicola* populations have shown varying compositions of companion symbionts, including *Arsenophonus*, *Pectobacterium*, *Serratia*, and *Wolbachia* alongside *Buchnera*, with variations among different populations (69). These observations, together with the distinct *Arsenophonus* origins among Cerataphidini species described above, indicate that companion symbiont associations in this tribe are highly dynamic, with frequent replacements occurring even between closely related species or within the same species. Moreover, in some Cerataphidini species, *Buchnera* has been replaced by yeast-like symbionts (70), further emphasizing the plasticity of symbiotic relationships in this lineage.

Such frequent replacements of companion symbionts are common features of dual-symbiotic systems. For example, in the Lachninae subfamily, *Serratia symbiotica* has been acquired by a common ancestor, establishing a dual-symbiotic system alongside *Buchnera* (67, 71). Within Lachninae, two distinct lineages of *Serratia symbiotica* have been described: one lineage, harbored by *Cinara tujafilina*, has a relatively large genome and is phylogenetically related to facultative *Serratia* in *A. pisum*, while another lineage, harbored by *Cinara cedri* and *Tuberolachnus salignus*, has a highly reduced genome (19, 20, 71, 72). Moreover, Lachninae species have undergone symbiont replacement from *Serratia* to *Erwinia*, *Sodalis*, and *Fukatsuia* symbionts (67, 73). These parallel evolutionary patterns demonstrate that companion symbionts can be frequently replaced even among closely related host species, while the ancient obligate symbiont *Buchnera* remains largely stable. Moreover, in the Auchenorrhyncha insects such as cicadas, spittlebugs, planthoppers, and leafhoppers, the ancient Bacteroidetes symbiont *Sulcia muelleri* has been maintained for over 270 million years, while its partner betaproteobacterial symbionts have been repeatedly replaced throughout evolutionary history (7, 9, 74–76). Similarly, in mealybugs with nested endosymbiosis, the inner gammaproteobacterial symbionts have been replaced multiple times while the outer *Tremblaya* symbionts remain stable (77). Together, these findings suggest that acquisition of new companion symbionts represents a common consequence to compensate for long-term genomic degradation in ancient obligate symbionts, thereby maintaining essential metabolic pathways critical for host survival.

## Conclusion

Our investigation of the subfamily Hormaphidinae reveals a complex evolutionary landscape of multi-partner symbioses, exemplified by the patterns observed in Figure 7. The tribe Cerataphidini demonstrates remarkable evolutionary dynamics, characterized by acquisition of a clade A *Arsenophonus* and extreme genome reduction in the primary symbiont *Buchnera* (∼0.4 Mbp) in the common ancestor, followed by frequent turnover of companion symbionts with diverse phylogenetic origins, and specialized bacteriocyte differentiation for newly acquired companion symbionts (Table 2). In contrast, the examined species of Hormaphidini and Nipponaphidini harbor only *Buchnera* with larger genomes, suggesting that the transition to dual-symbiosis occurred within Cerataphidini. Building on these findings, future studies examining a broader range of Hormaphidinae species will further resolve the evolutionary dynamics of dual-symbiosis across this subfamily.

**Table 2.**
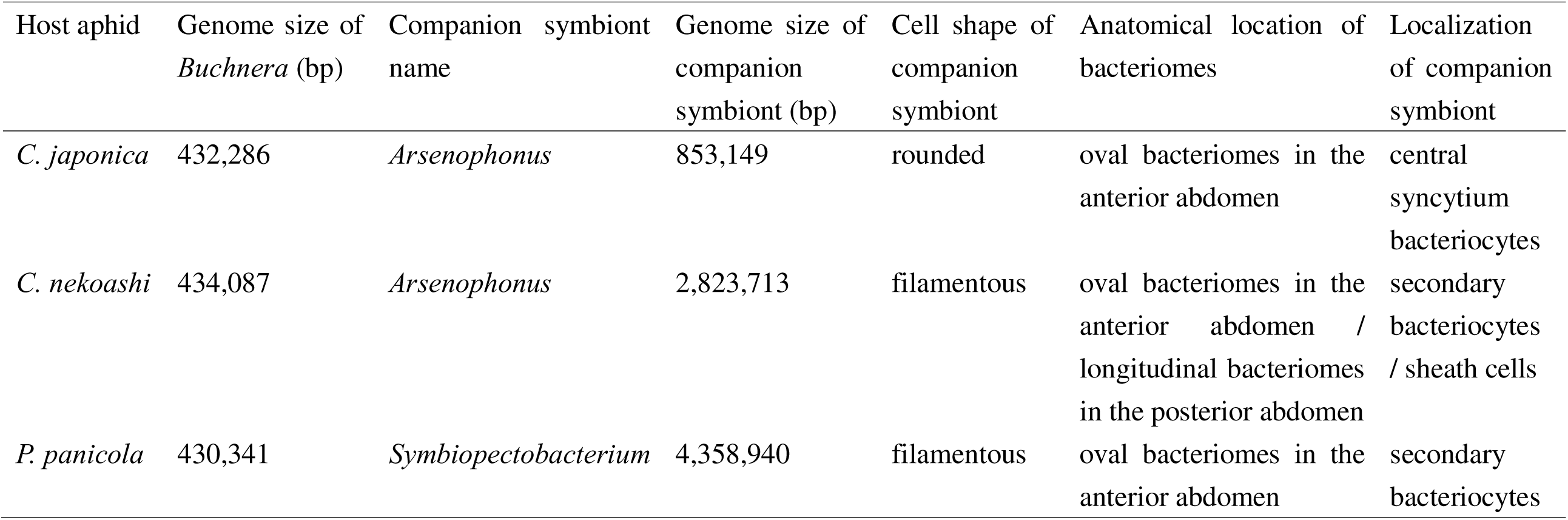
Comparison of symbiotic status among three Cerataphdini species.

## Materials and Methods

### Aphid collection and rearing

We collected eight Hormaphidinae species across three tribes: Hormaphidini, Nipponaphidini, and Cerataphidini. Aphid species were identified based on their morphological characteristics and host plants. The host plants and sampling locations are listed in Table S1. These aphid samples were preserved in 100% ethanol until 16S ribosomal RNA gene amplicon sequencing was performed.

We established isofemale strains of *C. nekoashi* and *P. panicola*. *C. nekoashi* was collected from the Japanese stiltgrass *Microstegium vimineum* in Okazaki, Aichi Prefecture, Japan (161 m above the sea level; 34°56’53.3”N, 137°13’14.1”E) in September 2022. *P. panicola* was collected from the wavyleaf basketgrass *Oplismenus undulatifolius* in Nagasaki, Nagasaki Prefecture, Japan (43 m above the sea level; 32°47’13.7”N, 129°51’00.3”E) in November 2020. Both *C. nekoashi* and *P. panicola* have been maintained in the asexual viviparous phase on their secondary host plants: *M. vimineum* for *C. nekoashi* and *O. undulatifolius* for *P. panicola*. The aphids are reared in an incubator under controlled conditions: 20 °C, 60% relative humidity, and a long-day photoperiod (16 hours light/8 hours dark cycle).

### 16S ribosomal RNA gene amplicon sequencing analysis

The genome DNA (gDNA) was extracted from eight Hormaphidinae species using a DNeasy Blood & Tissue Kit (Qiagen, Hilden, Germany) following the manufacturer’s instruction. Prior to DNA extraction, aphid samples were rinsed twice with fresh 70% ethanol. Each gDNA sample was derived from one to ten individuals. For samples with low DNA concentration, we performed ethanol precipitation with the Ethachinmate (Nippon Gene, Tokyo, Japan) to concentrate the gDNA.

The V3–V4 hypervariable regions of the 16S rRNA gene were amplified using either the primer set 341F (5′-CCTACGGGNGGCWGCAG-3′) and 785R (5′-GACTACHVGGGTATCTAATCC-3′) (25), or the Quick-16S™ Primer Set V3–V4 (Zymo Research, Irvine, CA, USA) (see Table S1 for primer information for each sample). Amplicon libraries were quantified with the KAPA SYBR FAST qPCR Kit (Kapa Biosystems, Wilmington, MA, USA) using the Applied Biosystems 7500 Real-Time PCR System (Applied Biosystems, Foster City, CA, USA), and sequenced on the Illumina MiSeq platform (Illumina, San Diego, CA, USA) to generate 250 bp paired-end reads, which yielded 777,518 raw reads. Primer sequences were trimmed from reads: 17 bp from the 5′ end of forward reads and 21 bp from the 5′ end of reverse reads for the 341F-785R primer set; 16 bp from the 5′ end of forward reads and 24 bp from the 5′ end of reverse reads for the Quick-16S™ Primer Set V3–V4. After quality filtering and removal of chimeric sequences using QIIME2 (version 2020.8) (26), 696,138 reads remained for analysis. Taxonomy assignment was performed against the Silva138 database (27). The raw Illumina sequencing data have been deposited in the DDBJ database under accession number DRR697964-DRR697984.

### Molecular phylogenetic analysis

Protein sequences of related bacteria were downloaded from NCBI (accession numbers in Table S2). Single-copy orthologous proteins were extracted using OrthoFinder (version 2.5.5) (28). For *Arsenophonus* symbionts detected by 16S rRNA gene amplicon sequencing but lacking genome assemblies, 16S rRNA gene sequences of related bacteria were additionally downloaded from NCBI to assess their phylogenetic positions (accession numbers in Table S2). Multiple sequence alignments were performed using MAFFT with the L-INS-i algorithm (version 7.520) (29). For protein sequences, gaps in the alignments were removed using TrimAL (version 1.4.rev15) with the “-gt 1.0” option (30). Trimmed protein sequences were then concatenated using our custom Python script. The best models were determined using ModelTest-NG (version 0.1.7) (31) as follows: CPREV+I+G4+F for protein phylogeny of *Buchnera* and *Arsenophonus*, TVM+I+G4 for 16S rRNA gene phylogeny of *Arsenophonus*, and JTT+I+G4+F for protein phylogeny of *Pectobacterium*. Maximum likelihood (ML) trees were reconstructed using RAxML-NG (version 1.2.0) (32) with the aforementioned models. Bootstrap values for ML phylogeny were obtained from 1,000 replicates. Phylogenetic trees were visualized using ggtree (version 3.12.0) (33). A multiple sequence alignment of the V3–V4 region of the 16S rRNA gene among *Arsenophonus* strains from Cerataphidini aphids was visualized using Jalview (version 2.11.2.0) (34).

### Genome sequencing

Genomic DNA was extracted from fresh isofemale strains of *C. nekoashi* and *P. panicola* maintained in our laboratory. Aphids were rinsed with fresh 70% ethanol with vigorous agitation to remove wax from the surface of their bodies, rinsed with Milli-Q water, and then blotted dry. To prepare for high molecular weight (HMW) DNA, whole-body samples of 50 individuals each of *C. nekoashi* and *P. panicola* were transferred to a mortar and gently ground into a fine powder with liquid nitrogen. Frozen powdery QIAGEN G2 buffer (Qiagen), generated by adding 2-mercaptoethanol to QIAGEN G2 buffer and spraying the mixture into liquid nitrogen in a glass beaker, was added to the samples and quickly blended. After allowing the mixture to thaw in a tube, RNaseA (Qiagen) and Proteinase K (Qiagen) were added, and the sample was incubated at 37°C for 2.0 h without agitation. The samples were centrifuged at 10,000 x g at 4°C for 20 min, and the supernatant was subjected to DNA extraction using a QIAGEN Genomic-tip 20/G column. Subsequent procedures followed the manufacturer’s instruction with minor modifications: the gDNA was eluted with 2 mL of Buffer QF, purified with AMPure XP beads (Beckman Coulter, Brea, CA, USA), and then eluted with 50-60 µL EB buffer. Note that the extracted gDNA included genomes derived from both aphids and their symbionts, i.e., hologenome. The quantity of extracted gDNA was measured using a Qubit dsDNA HS Assay Kit (Thermo Fisher Scientific, Waltham, MA, USA) and a Qubit 2.0 Fluorometer (Thermo Fisher Scientific). The quality of extracted gDNA was assessed using a Nanodrop ND-2000C (Thermo Fisher Scientific). The integrity of the HMW genome was assessed by a pulsed-field gel electrophoresis (PFGE) using a CHEF Mapper (Bio-Rad, Hercules, CA, USA). HMW genome was sheared to an average target size of 40-50 kbp in two cycles using speed settings of 29 and 30 on a Megaruptor^®^3 (Diagenode, Denville, NJ, USA). SMRTbell libraries for sequencing on the PacBio Sequel platform were constructed according to the manufacturer’s instruction (Preparing whole genome and metagenome libraries using SMRTbell^®^ prep kit 3.0 Procedure & checklist PN 102-166-600 APR2022) with minor modifications. Due to low DNA yields after nuclease treatment, samples were pooled and concentrated using SMRTbell cleanup beads prior to size selection using a BluePippin (Sage Scientific, Beverly, MA, USA). The size of the combined library was confirmed by PFGE using the CHEF Mapper as previously described for HMW genome evaluation. This SMRTbell library was sequenced in one SMRT cell (movie capture time, 20 h) on a Sequel IIe instrument.

### Genome assembly and annotation

Symbiont-derived HiFi reads were extracted by mapping to known *Buchnera*, *Arsenophonus*, and *Symbiopectobacterium* genome and protein sequences using BLASTN (version 2.13.0) (35) and DIAMOND (version 2.0.15) (36) with a BLASTX option. The extracted HiFi reads were deposited in the DDBJ database under accession number DRR697963 for *C. nekoashi* and DRR698109 for *P. panicola*.The extracted reads were used for metagenome assembly using metaMDBG (version 0.3) (37) with default options. *Buchnera*, *Arsenophonus*, and *Symbiopectobacterium* genomes were identified from the metagenome-assembled contigs based on DIAMOND searches against known symbionts and sequencing depth. To confirm circularity, HiFi reads were mapped to the symbiont genome sequences using minimap2 (version 2.24) (38), and the presence of overlapping reads at both ends was verified. We obtained closed circular sequences of *Buchnera* of *C. nekoashi* and *P. panicola*, *Arsenophonus* of *C. nekoashi*. The *Symbiopectobacterium* genome was assembled into two contigs, likely due to low read depth (∼8x) and long repeats at the 3’ end of the contig. The assembled genome sequences were deposited in the DDBJ database under the following accession numbers: AP041232-AP041234 for *Buchnera* of *C. nekoashi*, AP041230-AP041231 for *Arsenophonus* of *C. nekoashi*, AP041235-AP041237 for *Buchnera* of *P. panicola*, and AP041238-AP041239 for *Symbiopectobacterium* of *P. panicola*.

Gene predictions and annotations were performed using DFAST (version 1.3.6) (39). Although lengths of *ilvH* genes of *Buchnera* in *C. nekoashi* and *P. panicola* were less than ∼50% of the average length of their orthologs, we considered these as functional CDSs because of the retention of a functional N-terminal domain (40). The structural conservation of symbiont chromosomes was assessed by identifying syntenic regions within each of the following groups: four *Buchnera* strains and three *Arsenophonus* strains. Homology searches were performed using BLASTP (35) with the option “-outfmt 6”, and the results were visualized using pyGenomeViz (version 1.6.1; https://github.com/moshi4/pyGenomeViz). To identify shared and species-specific gene repertoires, we performed comparative genomic analysis using OrthoFinder (version 2.5.5) (28).

### Fluorescence *in situ* hybridization

For cryosections, third or fourth instar nymphs of *C. nekoashi* and *P. panicola* were embedded in Tissue-Tek^®^ O.C.T. Compound (Sakura Finetek Japan Co., Ltd, Tokyo, Japan) after removal of appendages to facilitate permeability. Samples were immediately frozen in liquid nitrogen and stored at −80°C until use. Frozen blocks were sectioned at 10 µm thickness using a Leica CM3050 S cryostat (Leica Biosystems, Nussloch, Germany), and sections were fixed with 4% paraformaldehyde dissolved in 1× phosphate buffered saline (PBS, pH 7.4) for 20 min, followed by washing with 0.3% Triton X-100 in 1× PBS (PBSTx).

For whole embryos, specimens were dissected from third or fourth instar nymphs and fixed with 4% paraformaldehyde dissolved in 1× PBS (pH 7.4) for 30 min. Fixed samples were dehydrated with 100% methanol for 30 min, then rehydrated through a graded series of PBSTx prior to the hybridization.

For both sample types, hybridization was performed in 20 mM Tris-HCl (pH 8.0), 0.9 M NaCl, 0.01% SDS, 30% (v/v) formamide containing DAPI (Dojindo Laboratories, Kumamoto, Japan) and 100 nM each of fluorescent-labeled oligonucleotide DNA probes for 2h. The following species-specific probes targeting 16S rRNA were used: Cy5_BucCn_16S_1267 (5’-Cy5-GTT CTC GCA AAT TCG CAT C-3’) for *Buchnera* of *C. nekoashi*, Cy3_ArsCn_16S_01846_1020 (5’-Cy3-CTG TCT CAG CGC TCC CGA AGG CAC TC-3’) for *Arsenophonus* of *C. nekoashi*, Cy5_BucPp_16S_53 (5’-Cy5-TTT CGC TGC CGC ACG ACT TGC-3’) for *Buchnera* of *P. panicola*, and Cy3_SymPp_16S_01534_837 (5’-Cy3-CCG GAA GCC ACG CCT CAA GGG-3’) for *Symbiopectobacterium* of *P. panicola*. After hybridization, excess probes were removed by washing with PBSTx. Specimens were mounted with VECTASHIELD (Vector Laboratories, Burlingame, CA, USA) and imaged using an Olympus FLUOVIEW FV1000 confocal laser scanning microscope (Olympus, Tokyo, Japan).

## Data availability

All sequencing data and assembled genome data have been deposited at the DDBJ under the bioProject ID: PRJDB35433. Accession numbers are written in the material and methods section. Original codes are available on GitHub (https://github.com/ShuntaYorimoto/25shunta_cnppsym_paper/tree/main).

## Acknowledgements

We appreciate Dr. Katsushi Yamaguchi, Dr. Asaka Akita, Hisayo Asao and Mika Ikeda in the Trans-Omics Facility, NIBB Trans-Scale Biology Center for technical support with Illumina and HiFi library preparation and sequencing. Computational resources were provided by the Data Integration and Analysis Facility, National Institute for Basic Biology and the Research Center for Computational Science, Okazaki, Japan (Project: NIBB, 24-IMS-C312, 25-IMS-C235). We thank three anonymous reviewers of an earlier version of this manuscript for their constructive comments, which helped improve the quality of this work.

## Author contributions

S. Y. and S. S. designed the research. S. Y., M. H., and T. N. collected aphid samples. S. Y. performed molecular biology experiments and microscopic analyses. S. Y. and S. S. performed next-generation sequencing and bioinformatics analyses. S. Y. and S. S. wrote the manuscript. All authors provided comments on revisions and approved the final manuscript.

## Supplemental item titles and legends

**Figure S1.** Phylogenetic analysis and sequence alignment of *Arsenophonus* based on the V3–V4 region of 16S rRNA genes (A) ML tree of *Arsenophonus* constructed using 435 variant patterns from 1,626 nucleotide positions in the 16S rRNA genes. *Proteus mirabilis* serves as an outgroup. Target symbionts in this study are highlighted in red. No bootstrap values are displayed in the tree as all nodes had support values below 70%. A scale bar represents 0.05 substitutions per site. (B) Multiple sequence alignment of 424 nucleotide positions from the V3–V4 region of 16S rRNA gene of *Arsenophonus* in *C. japonica*, *C. cerbera*, *C. nekoashi*, *C.* sp. B, *P. bambucicola*.

**Figure S2.** Synteny relationships of symbiont chromosomes (A) Genome-wide synteny among *Buchnera* chromosomes of *A. pisum*, *C. japonica*, *C. nekoashi*, and *P. panicola*. (B) Genome-wide synteny among *Arsenophonus* chromosomes of *C. japonica*, *C. nekoashi*, and *A. craccivora*. Chromosomes are represented as horizontal bars with host species names on the left and chromosome sizes below. Gray links indicate syntenic regions in the same orientation; red links indicate inverted syntenic regions. Color intensity reflects sequence identity (30–100%).

**Table S1.** Bacterial symbiont abundance and sampling information for Hormaphidinae species

**Table S2.** 16S rRNA gene and protein sequences used for phylogenetic analyses and alignments

**Table S3.** Genes commonly lost in *Buchnera* genomes of three Cerataphidini aphids compared to that of *A. pisum*

