## Supplemental Figure for "Evolution of multi-partner symbiotic systems in the tribe Cerataphidini: genome reduction of Buchnera and frequent turnover of companion symbionts"

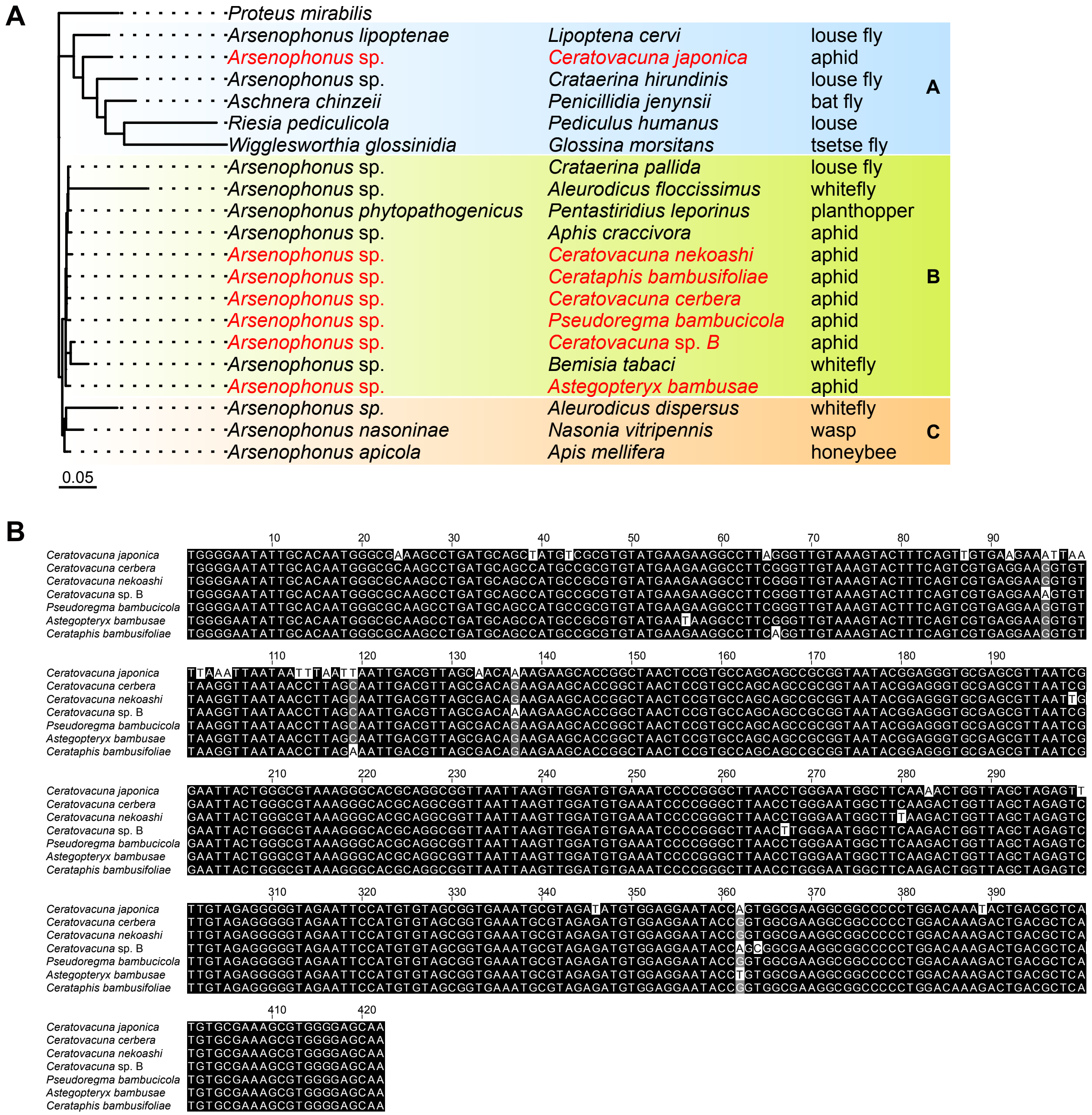


Figure S1. Phylogenetic analysis and sequence alignment of *Arsenophonus* based on the V3–V4 region of 16S rRNA genes

(A) ML tree of *Arsenophonus* constructed using 435 variant patterns from 1,626 nucleotide positions in the 16S rRNA genes. *Proteus mirabilis* serves as an outgroup. Target symbionts in this study are highlighted in red. No bootstrap values are displayed in the tree as all nodes had support values below 70%. A scale bar represents 0.05 substitutions per site. (B) Multiple sequence alignment of 424 nucleotide positions from the V3–V4 region of 16S rRNA gene of *Arsenophonus* in *C. japonica*, *C. cerbera*, *C. nekoashi*, *C.* sp. B, *P. bambucicola*.

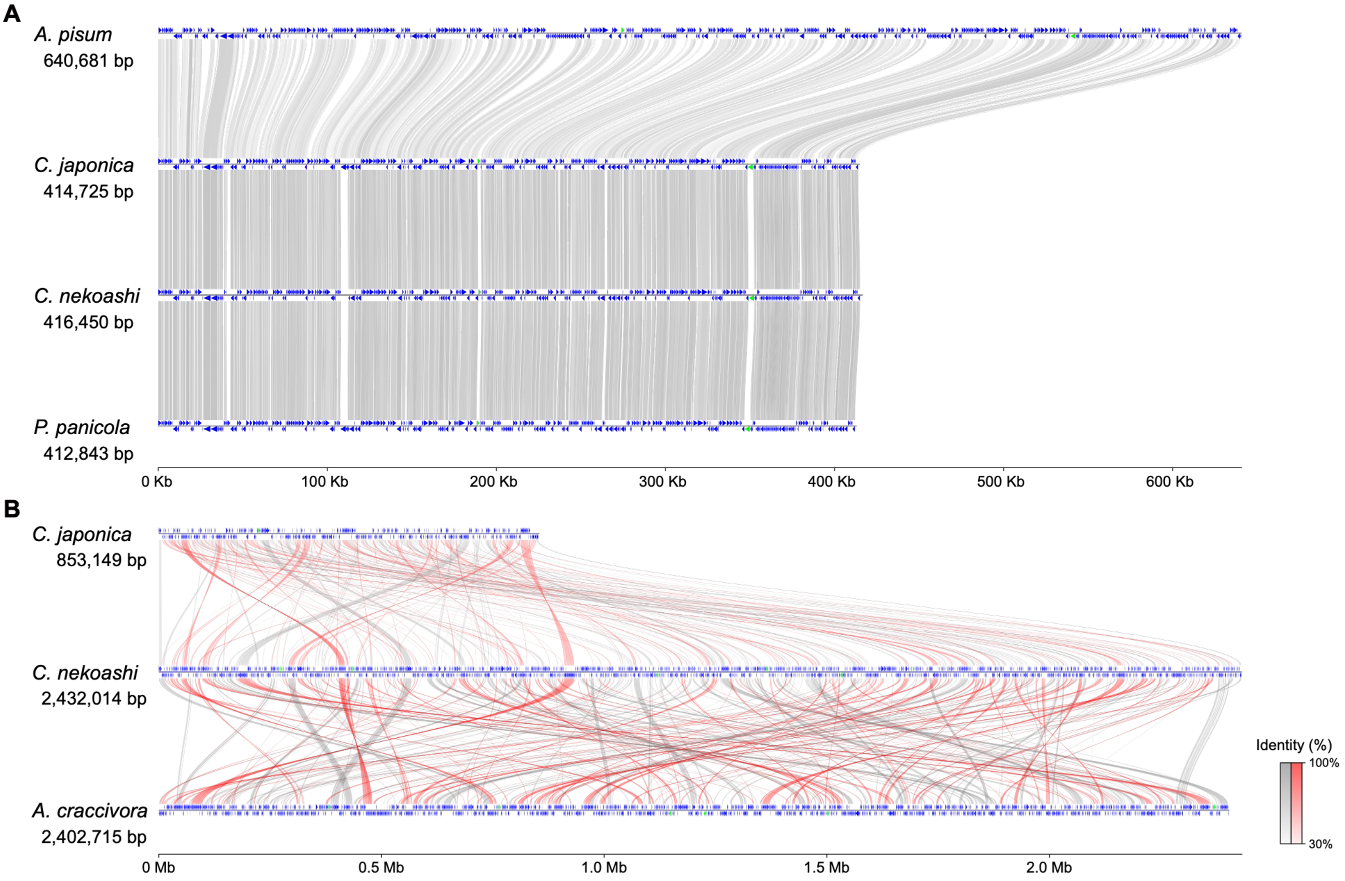


Figure S2. Synteny relationships of symbiont chromosomes

(A) Genome-wide synteny among *Buchnera* chromosomes of *A. pisum*, *C. japonica*, *C. nekoashi*, and *P. panicola*. (B) Genome-wide synteny among *Arsenophonus* chromosomes of *C. japonica*, *C. nekoashi*, and *A. craccivora*. Chromosomes are represented as horizontal bars with host species names on the left and chromosome sizes below. Gray links indicate syntenic regions in the same orientation; red links indicate inverted syntenic regions. Color intensity reflects sequence identity (30–100%).
